# The CONDOR pipeline for simultaneous knockdown of multiple genes identifies RBBP-5 as a germ cell reprogramming barrier in *C. elegans*

**DOI:** 10.1101/2020.09.01.276972

**Authors:** Marlon Kazmierczak, Carlota Farré i Díaz, Andreas Ofenbauer, Baris Tursun

**Affiliations:** Berlin Institute of Medical Systems Biology; Max Delbrück Center for Molecular Medicine in the Helmholtz Association, 13125 Berlin, Germany

**Author notes:** **Correspondence to:** (BT).

**Keywords:** RNA interference, Double RNAi, Genetic Screen, Conjugation, Reprogramming, Epigenetics, *C. elegans*

## Abstract

Multiple gene activities control complex biological processes such as cell fate specification during development and cellular reprogramming. Investigating the manifold gene functions in biological systems requires also simultaneous depletion of two or more gene activities. RNA interference-mediated knockdown (RNAi) is commonly used in *C. elegans* to assess essential genes, which otherwise lead to lethality or developmental arrest upon full knockout. RNAi application is straightforward by feeding worms with RNAi plasmid-containing bacteria. However, the general approach of mixing bacterial RNAi clones to deplete two genes simultaneously often yields poor results. To address this issue, we developed a bacterial conjugation-mediated double RNAi technique ‘CONDOR’. It allows combining RNAi bacteria for robust double RNAi with high-throughput. To demonstrate the power of CONDOR for large scale double RNAi screens we conjugated RNAi against the histone chaperone gene *lin-53* with more than 700 other chromatin factor genes. Thereby, we identified the Set1/MLL methyltransferase complex member RBBP-5 as a novel germ cell reprogramming barrier. Our findings demonstrate that CONDOR increases efficiency and versatility of RNAi screens to examine interconnected biological processes in *C. elegans* with high-throughput.

## INTRODUCTION

Most biological processes such as development, cell fate specification, aging, and behavior are controlled by the activity of multiple genes. One approach to investigate the implication of genes in regulating such complex processes is their inactivation to assess of related perturbation phenotypes (Boutros and Ahringer, 2008).

Reverse genetics by RNAi is an essential tool to at least partially inactivate genes in the nematode *C. elegans*, which has been used as a powerful model organism to reveal highly conserved molecular mechanisms and gene regulatory pathways (Dudley and Goldstein, 2005; Markaki and Tavernarakis, 2020). To perform RNAi in *C. elegans*, animals are fed individual *E. coli* bacterial strains producing dsRNA against only one specific gene (Conte et al., 2015; Kamath et al., 2003) (Fig. 1A). RNAi causes a partial knockdown allowing the investigation of genes, which would cause early developmental arrest, sterility, or even lethality when fully depleted. This RNAi feature is an important benefit compared to genetic screens based on mutagenesis or gene editing. Mutagenizing chemical compounds or CRISPR/Cas9 often lead to a full gene knockout, and hence, reduce the possibility to study essential genes during biological processes post-embryonically or in adult animals (Boutros and Ahringer, 2008).

**Figure 1:**
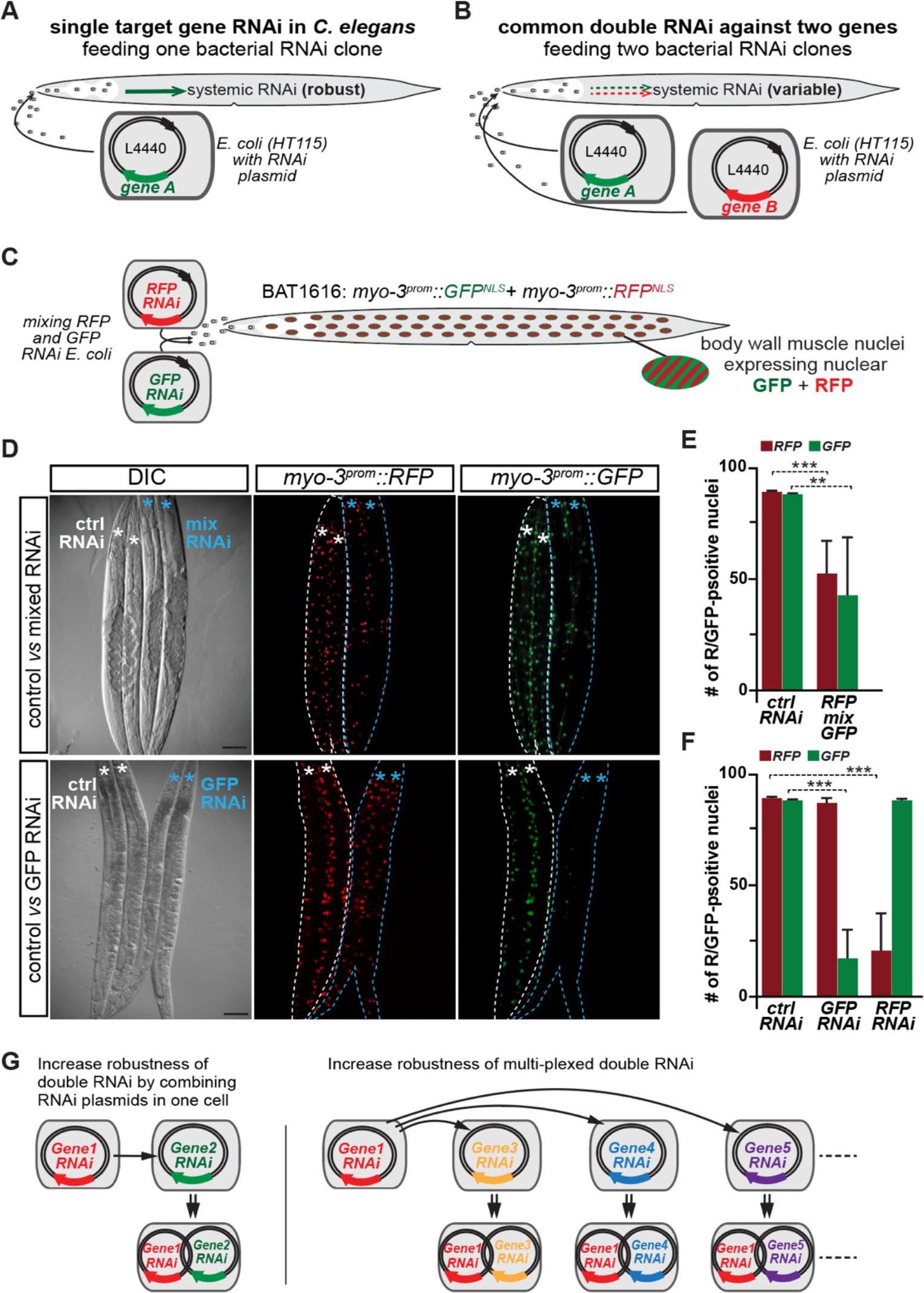
Double RNAi in C. elegans by feeding mixed RNAi bacteria. (A) RNAi in *C. elegans* is straightforward by feeding dsRNA-producing *E. coli (HT115* strain). dsRNA against the target gene is produced from the L4440 RNAi plasmid. (B) Double RNAi to knockdown two genes simultaneously by mixing two RNAi bacteria clones. (C) Illustration of transgenic BAT1616 worms expressing RFP and GFP in nuclei of muscles. Using the *myo-3* promoter 95 body wall muscle nuclei in hermaphrodites are labelled. (D) Representative pictures of DIC and fluorescent signals of BAT1616 fed with RNAi bacteria against RFP and GFP either mixed or individually. Asterisks label pharynx of simultaneously imaged animals. Scale bars are 20 μm. (E and F) Quantification of muscle nuclei number with depleted GFP or RFP signals. Statistics: unpaired t-test; ***p < 0,0001; **p < 0,001. n= 200. Error bars represent SEM (G) Illustration of proposition to increase robustness of double RNAi by combining two RNAi plasmids in bacterial cells.

Simultaneous RNAi-mediated knockdown of two genes in *C. elegans* is generally applied by mixing two bacterial strains that contains specific dsRNA-producing plasmids targeting an individual gene (Kamath et al., 2001) (Fig. 1B). However, this approach is not reliable and often yields inefficient knockdown of both genes (Gouda et al., 2010). This inefficiency can be overcome by generating a single plasmid producing both dsRNAs against the targeted genes (Gouda et al., 2010). While ‘stitching’ target genes together on one RNAi plasmid mediates robust double RNAi, it is not feasible for large scale screens, as it would require high-throughput plasmid cloning.

In order to reconcile double RNAi robustness with high-throughput screening, we developed a <u>CON</u>jugation-mediated <u>DO</u>uble <u>R</u>NAi technique, which we term ‘CONDOR’. CONDOR generates double RNAi bacteria clones in high-throughput and significantly reduces the amount of time and reagents compared to plasmid cloning. At the same time CONDOR provides simultaneous knockdown of a large set of two-gene combinations in a robust manner.

To examine the efficiency of CONDOR for large-scale screening, we investigated the knockdown of around 700 chromatin factors in combination with RNAi against the histone chaperone LIN-53 in *C. elegans*. LIN-53 was previously identified to prevent in conjunction with the chromatin silencer PRC2 transcription factor-induced (TF) conversion of germ cells into neuron-like cells (Patel et al., 2012; Seelk et al., 2016; Tursun et al., 2011). RNAi against *lin-53* alone allows efficient germ cell conversion to glutamatergic neurons (termed ASE) by the TF CHE-1. In contrast, only limited conversion to GABAergic motor neurons by the Pitx-type homeodomain TF UNC-30 could be observed in LIN-53-depleted animals. We hypothesized that depletion of additional chromatin regulators together with *lin-53* may increase germ cell reprogramming to GABAergic neurons. Indeed, our CONDOR screen revealed that co-depletion of the Set1/MLL methyltransferase complex member RBBP-5 together with LIN-53 significantly increased germ cell reprogramming to GABAergic neurons.

Chromatin factors have been identified in *C. elegans* and mammals as cellular reprogramming barriers and possible interplay of epigenetic mechanisms remains to be an important research aspect of safeguarding cell fates. CONDOR provides the multiplexed interrogation of combinatorial gene knockdowns for investigating such aspects, but can be also applied in the context of other biological phenomena. Genes may act in distinct or similar pathways with identical or converging physiological functions. Hence it is important to study their relationship in order to better understand underlying mechanisms of biological processes such as development and cell fate specification, which may also be relevant for addressing open questions in biomedical research.

## RESULTS

### Double RNAi by mixing bacterial strains is inefficient

In order to assess the degree of double RNAi robustness we generated the strain BAT1616, which expresses red fluorescent protein (RFP) as well as green fluorescent protein (GFP) in muscles using the *myo-3* promoter (Fig. 1C). For easy assessment of fluorescence signal intensities both fluorescent proteins are localized to the nuclei of all 95 body wall muscles (Fig. 1C-D). To simultaneously deplete RFP and GFP signals in the muscle nuclei we applied a 1:1 mix of RNAi bacterial clones, based the ‘Ahringer’ RNAi library *HT115 E. coli* strains, each containing RNAi plasmids against *RFP* and *GFP* (Fig. 1C-F) (Supp. Fig. 1). Around 50% of muscle nuclei lost RFP and GFP signals in F1 RNAi animals (Fig. 1C-F). In contrast, feeding *HTT115 E. coli* with GFP or RFP RNAi-plasmids individually reduced GFP and RFP signals more efficiently in approximately 75% of muscle nuclei, respectively (Fig. 1F). This outcome confirmed that mixing RNAi bacteria attenuates knockdown efficiency of individual genes as previously reported (Gouda et al., 2010).

To solve the issue of decreased RNAi efficiency upon mixing different RNAi bacteria, we assumed that combining two different RNAi plasmids in the same cell may increase robustness and efficiency of double RNAi (Fig. 1G).

### Bacterial conjugation to combine RNAi plasmids

Generating hundreds or thousands of new plasmids as described previously (Gouda et al., 2010) to combine a target gene of interest with a set of other targets is not feasible for large-scale double RNAi screens.

In order to develop a pipeline that allows combining RNAi plasmids with high-throughput, which provides robust double RNAi knockdowns, we sought for a method that consumes low amount of time and reagents. One such approach is bacterial conjugation, which allows the transfer of plasmids with an *origin of transfer* (*oriT*) among bacterial cells. Competence for bacterial conjugation requires presence of the fertility factor, also termed F-plasmid, which contains several genes of the *tra* locus for the formation of a pilus appendage. Bacteria with the F-plasmid are denoted as F^+^ (donor) and connect via the pilus to F^-^ bacteria (recipient) to transfer plasmids or other genetic material containing an *oriT* to the recipient (Supp. Fig. 2).

To adopt bacterial conjugation for combining RNAi plasmids, we made the F-plasmid *pRK24* (Meyer et al., 1979) and the RNAi plasmid *L4440* to be compatible with each other. We first replaced the Ampicillin resistance (AmpR) of *pRK24* with Kanamycin resistance (KanR) because *L4440* used in the standard ‘Ahringer’ *C. elegans* RNAi library (Kamath et al., 2003) already carries AmpR. To exchange AmpR with KanR we used recombineering, as previously described, due to the extensive size of *pRK24* (Fig. 2A) (Tursun et al., 2009). The generation of a selectable ‘donor’ RNAi plasmid, which can be transferred by conjugation, needed the addition of the *oriT* and replacement of AmpR with Chloramphenicol (CamR) resistance. This allows selection for presence of the transferred RNAi plasmid together with the resident AmpR-containing *L4440* RNAi plasmid after conjugation. We termed the newly generated donor plasmid ‘*LoriT*’, which is basically L4440 carrying *oriT* and CamR instead AmpR (Fig. 2B). Additionally, we figured that maintaining a large episome such as the F-plasmid requires stable conditions in bacteria - optimally preventing recombination events. Therefore, we used the *E. coli* strain *EPI300*, which is deficient of recombinases and has proven to maintain large fosmids in a stable manner (Fig. 2C) (Tursun et al., 2009). After cloning the target gene into *LoriT*, it is transformed to F^+^ *EPI300* bacteria (contain *pRK24-Kan*). This creates the donor strain that is ready to be conjugated with the receiving ‘Ahringer’ RNAi bacteria (in *HT115 E.coli* strain) for combining the target gene of interest with any other target gene for double RNAi (Fig. 2C).

**Figure 2:**
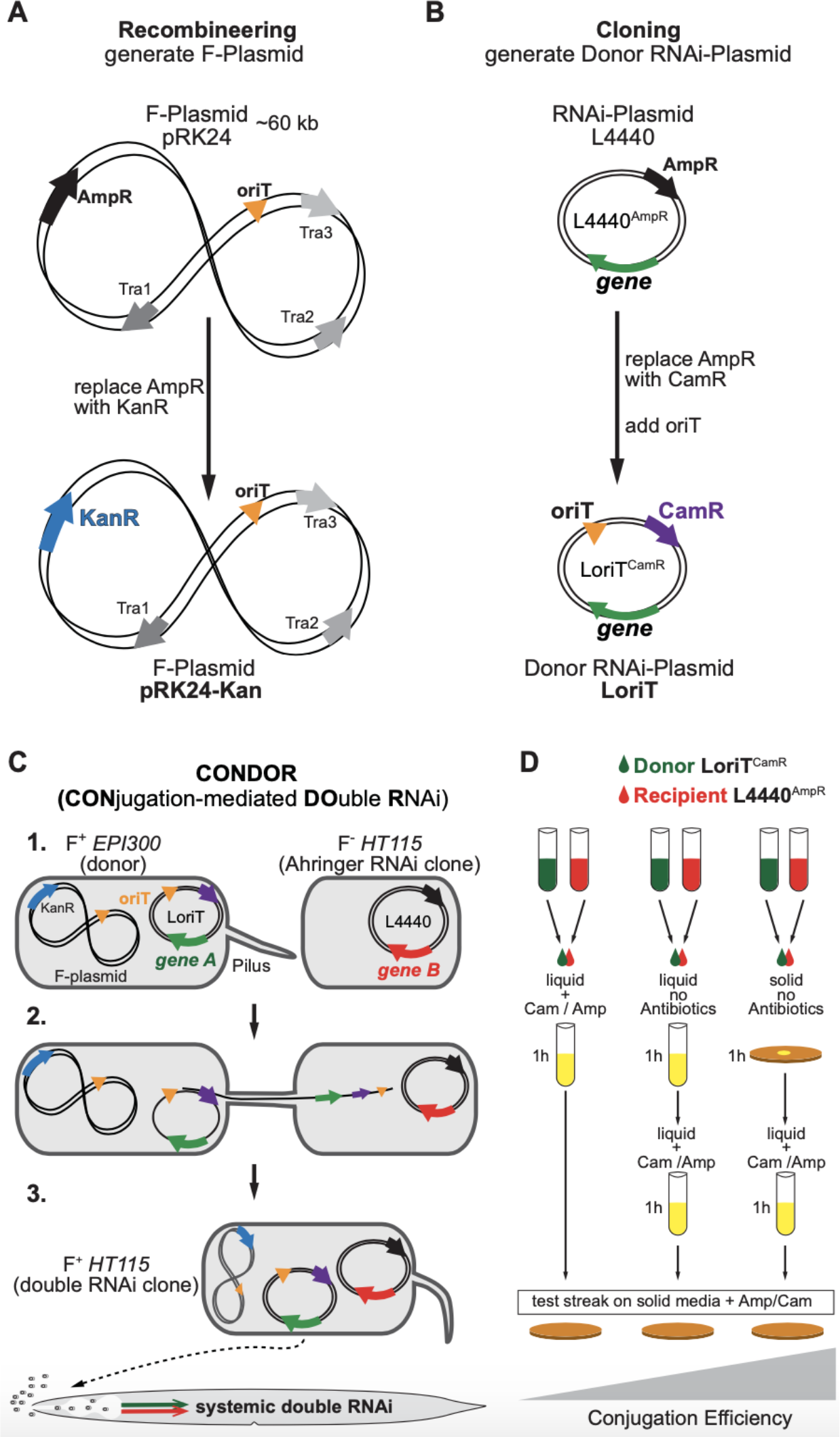
Creating a double RNAi system by bacterial conjugation. (A) The F-plasmid (fertility factor) encodes for components of the conjugation machinery to transfer *oriT*-containing genetic material. Recombineering was used to replace Ampicillin resistance (AmpR) with Kanamycin resistance (KanR) to allow combination with RNAi plasmids. (B) *LoriT* plasmid: we equipped the *L4440* RNAi plasmid (used for clones of the Ahringer RNAi library) with *oriT* and exchanged AmpR with Chloramphenicol resistance (CamR). (C) *pRK24-KanR-containing EPI300 E. coli* are F^+^ and can conjugate with *HT115* RNAi bacteria clones of the Ahringer RNAi library, which are F^-^. Conjugated bacteria are selected based on CamR / AmpR. (D) Different conjugation procedures were evaluated to find the most efficient transfer of *LoriT* to recipient RNAi bacteria. For detailed description and results see Supplemental Figure 3.

Next, we tested which conjugation procedure yields the most efficient transfer of the target gene-containing *LoriT* (Fig. 2D, Supp. Fig. 3A-I). By analyzing a number of variations including antibiotics at different steps and performing conjugation in liquid versus on solid media we determined the most efficient procedure (Supp. Fig. 3A-I). Combining a 5:1 ratio of donor: recipient bacterial culture on solid LB agar for 1h with subsequent selection (Cam / Amp in liquid for 1h) yielded 100% conjugation efficiency (Fig. 2D, Supp. Fig. 3A-I).

Overall, our adaption of the bacterial conjugation system to combine RNAi plasmids in bacteria is robust and straightforward to generate a large set of bacterial cells simultaneously producing dsRNA against two target genes. We termed our new technique CONDOR, which stands for <u>CON</u>jugation-mediated <u>DO</u>uble <u>R</u>NAi.

### Robust knockdown of two GFP and RFP in muscles by CONDOR

To demonstrate that CONDOR provides efficient knockdown of two genes simultaneously we co-depleted GFP and RFP expressed in muscle nuclei as described before. Additionally, we generated a *LoriT-GFP* RNAi plasmid and transformed into the F^+^ *EPI300 E. coli* strain (containing *pRK24-KanR*) (Fig. 3A). Subsequent conjugation with standard *HT115 E. coli* containing *L4440-RFP* RNAi plasmid generated bacterial cells producing dsRNA against both GFP and RFP (*GFP_CON_RFP*) (Fig. 3A-B). To assess and compare knockdown efficiencies, the conjugated bacteria and mixed GFP+RFP RNAi bacteria were used for double RNAi in BAT1616 worms (Fig. 3B-C). A similar result for mixed RNAi bacteria was observed as described before, where around 50% of nuclei showed simultaneous GFP/RFP signal depletion (Fig. 1D-E; Fig. 3C-D). In contrast, the conjugated bacteria simultaneously depleted GFP und RFP signals in more than 90% of the cells.

**Figure 3:**
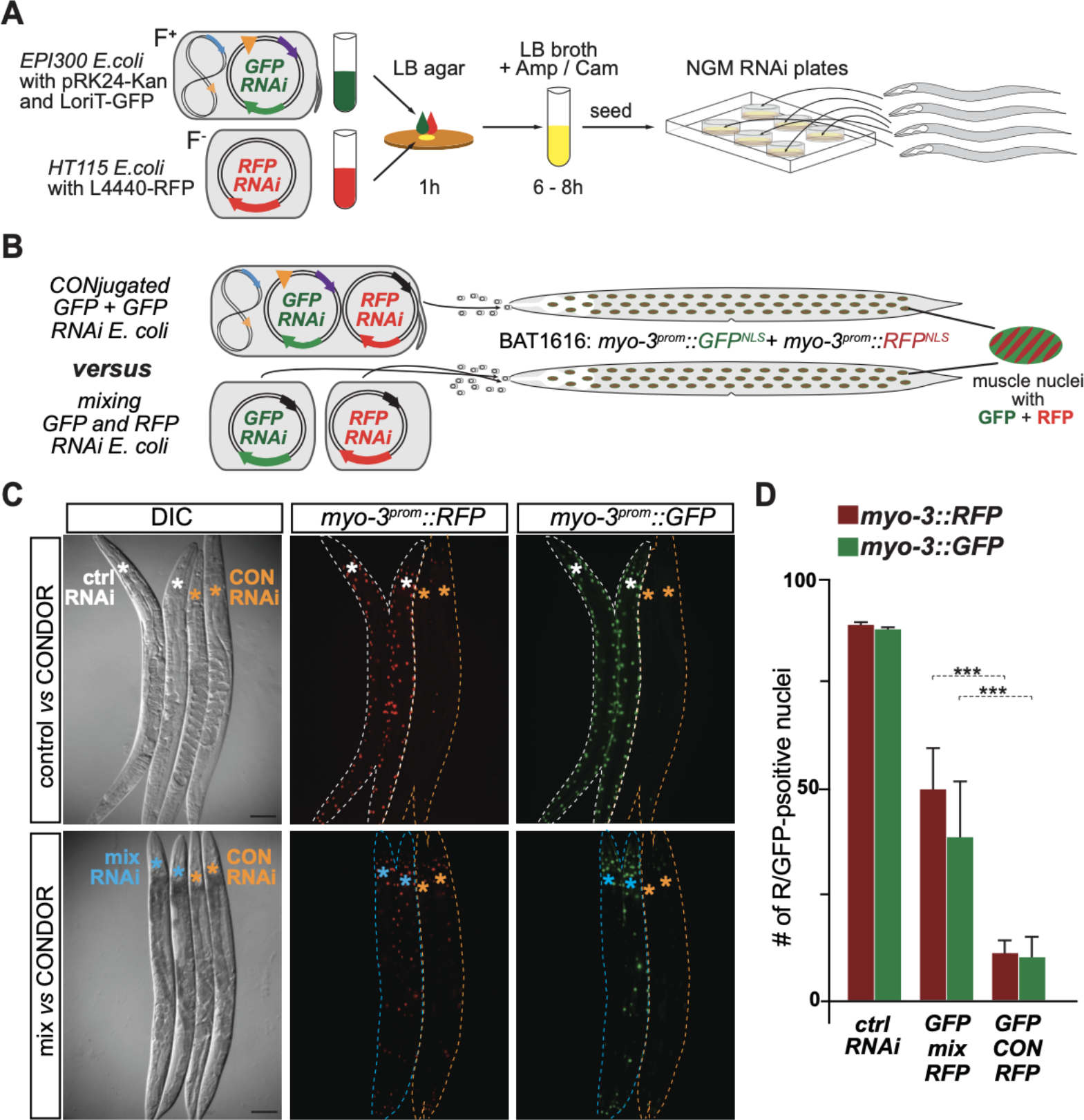
CONDOR knockdown of nuclear GFP and RFP in muscles. (A) Schematic illustration of CONDOR to generate *GFP* and *RFP* double RNAi bacteria. (B) Double RNAi against GFP and RFP in muscle nuclei of BAT1616 by CONDOR versus mixing individual RNAi bacteria. (C) Representative pictures of DIC and fluorescent signals of BAT1616 fed with RNAi bacteria against *RFP* and *GFP* either mixed or conjugated. Asterisks label pharynx of simultaneously imaged animals. Scale bars are 20 μm. (D) Quantification of muscle nuclei number with depleted GFP or RFP signals. CONDOR is depleting GFP and RFP more efficiently than mixing RNAi bacteria. Statistics: unpaired t-test; ***p < 0,0001; n= 120. Error bars represent SEM.

The outcome of testing BAT1616 worms for double RNAi against GFP and RFP indicates a very robust double RNAi knockdown by combining RNAi plasmids in one cell via conjugation using CONDOR.

### CONDOR is a robust double RNAi approach also in other systems

To assess the efficiency of CONDOR in targeting endogenous genes, we decided to target *oma-1* and *oma-2*, which are redundantly required for oocyte maturation (Detwiler et al., 2001). Double mutants lacking *oma-1* and *oma-2* are sterile due to immature oocytes, which accumulate in the gonads. In contrast, animals missing only *oma-1* or *oma-2* are fertile (Detwiler et al., 2001) (Fig. 4A). Wild-type worms were subjected to F1 double RNAi targeting *oma-1* and *oma-2* either by mixing the individual *oma-1* and *oma-2 RNAi* bacteria, or by using conjugated *oma-1_CON_oma-2* RNAi bacteria (Fig. 4B-D). In this context, we also tested whether different double RNAi clones generated by conjugation are equally effective. While mixed RNAi caused an arrest of oocyte maturation in around 25% of animals, three different conjugation-derived double RNAi bacteria clones against *oma-1_CON_oma-2* caused around 60% sterility in a reproducible manner (Fig. 4C-D). Notably, animals showed the characteristic *‘oma’* phenotype, which leads to accumulation of immature oocytes in the gonad, indicating that sterility was indeed caused due to depletion of *oma-1* and *oma-2* (Fig. 4D).

**Figure 4:**
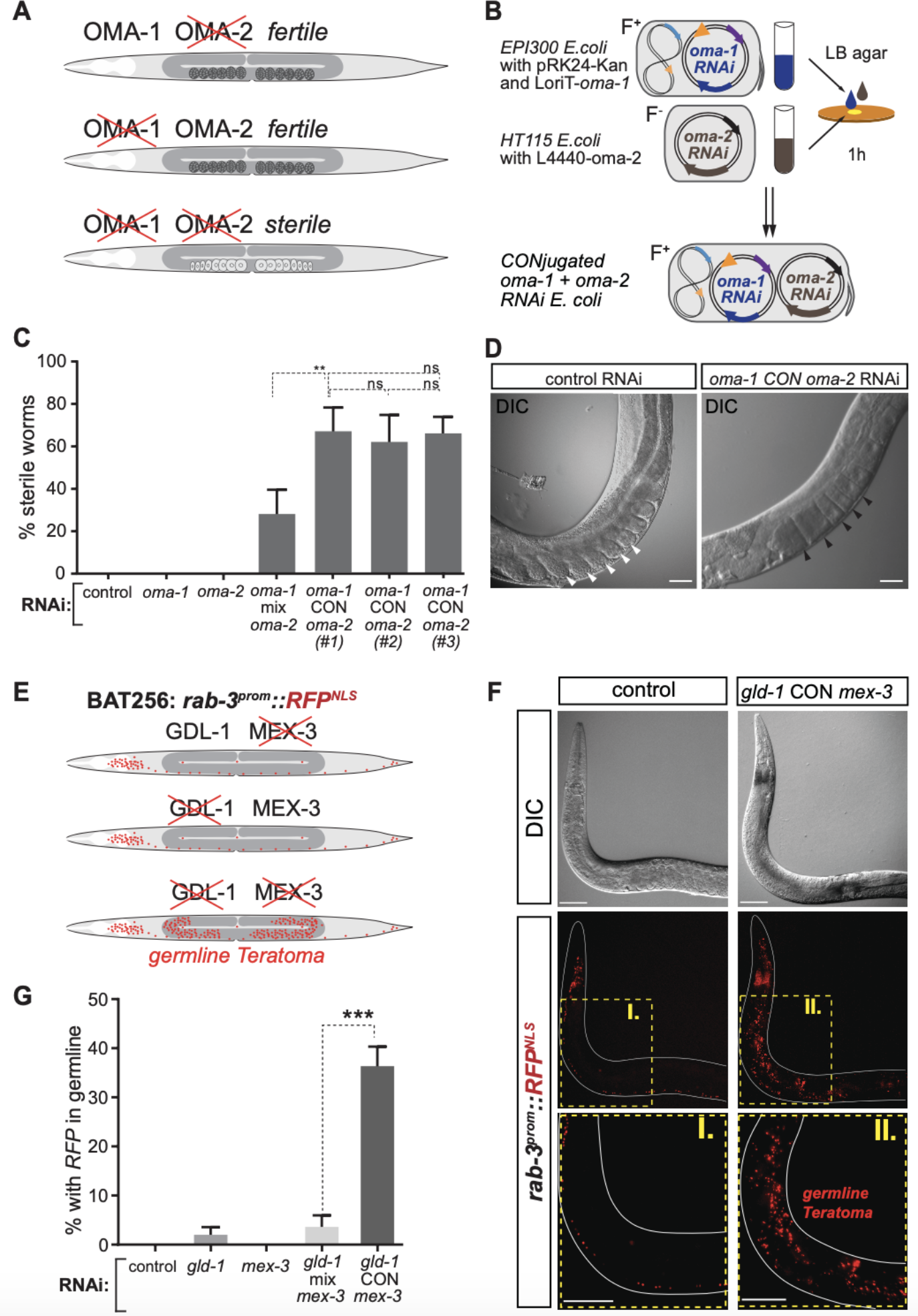
CONDOR knockdown of two endogenous genes. (A) Double depletion of *oma-1* and *oma-2* causes sterility due to immature oocytes. (B) Schematic illustration of CONDOR to generate *oma-1*_CON_ *oma-2* double RNAi bacteria. (C) Three independent *oma-1*_CON_ *oma-2* bacteria clones were tested and compared to mixing RNAi bacteria against *oma-1* and *oma-2*. Quantification of sterile animals displays higher efficiency of CONDOR for all three tested *oma-1*_CON_ *oma-2* clones compared to mixed RNAi bacteria. Statistics: unpaired t-test; **p < 0,001; n= 140, ns = not significant. Error bars represent SEM. (D) Representative DIC pictures of gonad region of control animals and *oma-1*_CON_ *oma-2* double RNAi treated animals. White arrow heads depict embryos, black arrow heads indicate accumulation of immature oocytes. Scale bars are 10 μm. (E) Double depletion of *gld-1* and *mex-3* leads to teratoma formation in the germline. (F) Representative fluorescence signal pictures of animals expressing the pan-neuronal reporter *rab-3::RFP^NLS^*. Double RNAi using CONDOR against *gld-1* and *mex-3* leads to teratoma formation visualized by the expression of neuronal RFP signals in the germline. Scale bars are 10 μm. (G) Quantification of teratoma formation confirms significantly increased induction of teratoma formation upon feeding with *gld-1 _CON_mex-3* bacteria compared to mixed RNAi bacteria against *gld-1* and *mex-3*. Statistics: unpaired t-test; ***p < 0,0001; n= 100. Error bars represent SEM.

Furthermore, we assessed teratoma formation of germ cells upon depletion of the translational regulators GLD-1 and MEX-3. Mutants carrying both mutations *gld-1*(*q485*) and *mex-3*(*or20*) alleles have been shown to develop teratomas in their germline that can be visualized based on expression of pan-neuronal reporters (Ciosk et al., 2006). We used worms, which express RFP in neuronal nuclei under the control of the pan-neuronal *rab-3* gene promoter (Fig. 4 E-F). Using CONDOR to target *gld-1* and *mex-3* simultaneously caused significantly more animals (35%) with germline teratomas than mixing RNAi bacteria against both genes (5%) (Fig. 4F-G). Additionally, we also targeted the 26S-Proteasome subunit genes *rpn-10* and *rpn-12*, which cause synthetic lethality when co-depleted (Takahashi et al., 2002). While CONDOR-mediated simultaneous knockdown of *rpn-10* and *rpn-12* reduced survival by around 50%, only 25% of the animals fed with mixed *rpn-10* and *rpn-12* RNAi bacteria died (Supp. Fig. 4).

Overall, our results provide evidence that CONDOR is a highly robust technique for simultaneous knockdown of endogenous genes by double RNAi.

### Epigenetic barriers of germ cell reprogramming to neuron-like cells

It was previously discovered that the epigenetic factor LIN-53, which can directly bind to histones, acts as a reprogramming barrier in the germline. RNAi against *lin-53* allows germ cell reprogramming to defined types of neuronal cells upon overexpression of specific transcription factors (TF). Overexpression of the Zn-finger TF CHE-1 induces conversion to glutamatergic ASE neuron-like cells labelled by expression of the ASE neuron-specific reporter *gcy-5::GFP* (Fig. 5A). The Pitx-type homeodomain TF UNC-30 is required for the specification of GABAergic motor neurons. Consequently, its overexpression in *lin-53* RNAi animals induces the GABA fate marker *unc-25::GFP* in germ cells (Fig. 5B). However, the induction of *unc-25::GFP* by UNC-30 is less efficient than *gcy-5::GFP* induction by CHE-1 (Fig. 5C). We speculated that this discrepancy may be due to additional epigenetic barriers that limit ectopic induction of the GABAergic motor neuron fate (Fig. D).

**Figure 5:**
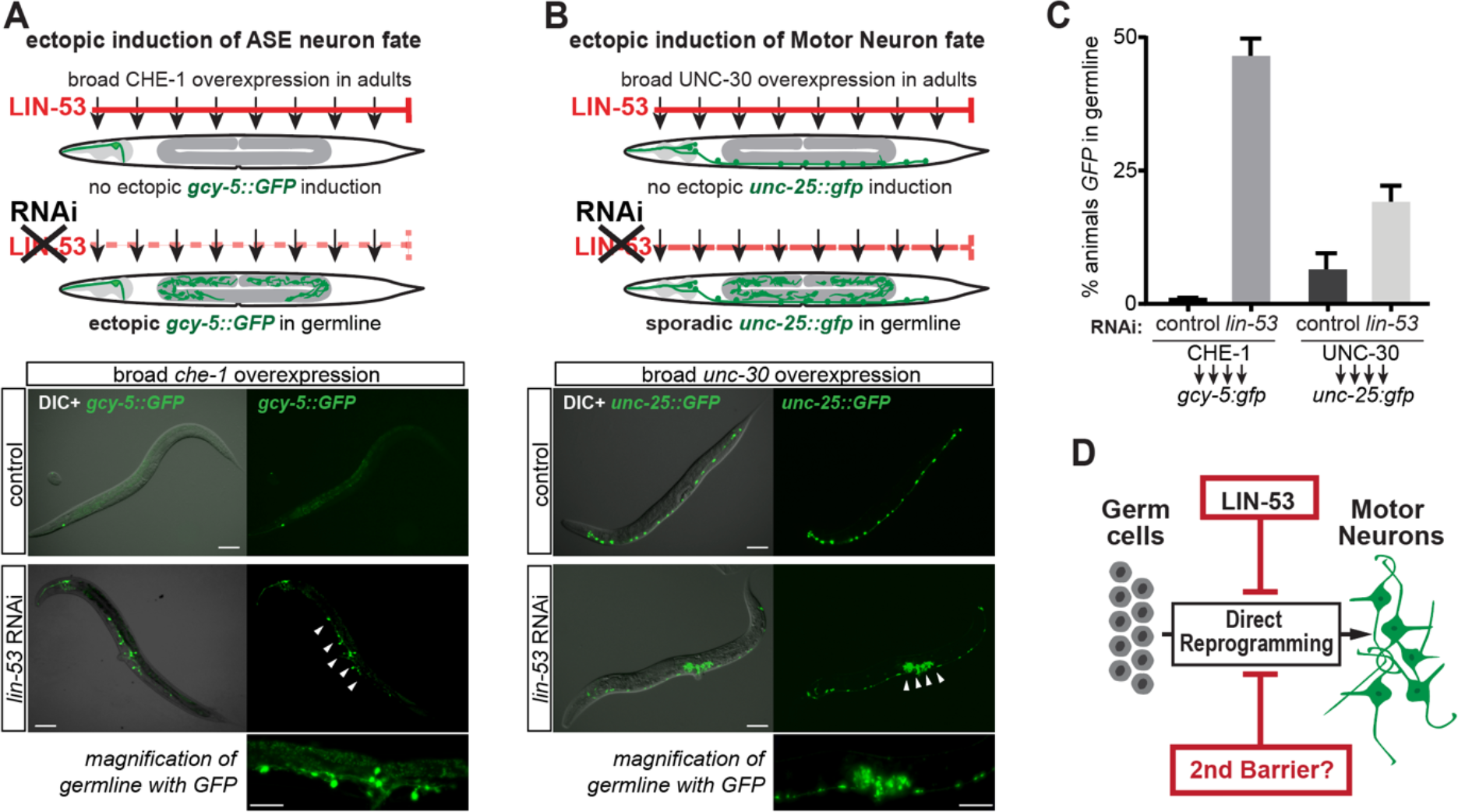
Epigenetic barriers of germ cell to neuron reprogramming in *C. elegans*. (A) Schematic illustration of transgenic animals expressing the glutamatergic ASE neuron fate marker *gcy-5::GFP* and allowing heat-shock-inducible broad CHE-1 overexpression. DIC / GFP pictures of animals with germ cells reprogrammed to ASE neurons upon depletion of the histone chaperone LIN-53 and broad overexpression of CHE-1. White arrow heads indicate germline with reprogrammed cells (this area is magnified below). (B) Schematic illustration of transgenic animals expressing the GABAergic motor neuron fate marker *unc-25::GFP* and allowing heat-shock-inducible broad UNC-30 overexpression. DIC / GFP signal pictures of animals with germ cells reprogrammed to GABAergic neurons upon depletion of LIN-53 and broad overexpression of UNC-30. White arrow heads indicate germline with reprogrammed cells (area is shown in magnification below). (C) Quantification of germ cell to neuron reprogramming by CHE-1 and UNC-30 upon *lin-53* RNAi. Induction of the GABA fate marker by UNC-30 is less efficient. Error bars represent SEM. (D) A second barrier may decrease germ cell to GABAergic motor neuron conversion.

The task to screen for a putative ‘2^nd^ barrier’ by co-depleting *lin-53* with other chromatin regulators provided an attractive test case to perform CONDOR for a large set of double RNAi knockdowns.

### CONDOR identifies RBBP-5 as a novel reprogramming barrier

To identify additional epigenetic regulators, which may be involved in limiting the conversion of germ cells to GABAergic neurons, we decided to conjugate the *LoriT-lin-53* plasmid with around 700 other RNAi clones that target chromatin-related genes based on our previously described chromatin RNAi library (Hajduskova et al., 2019) (Fig. 6A).

**Figure 6:**
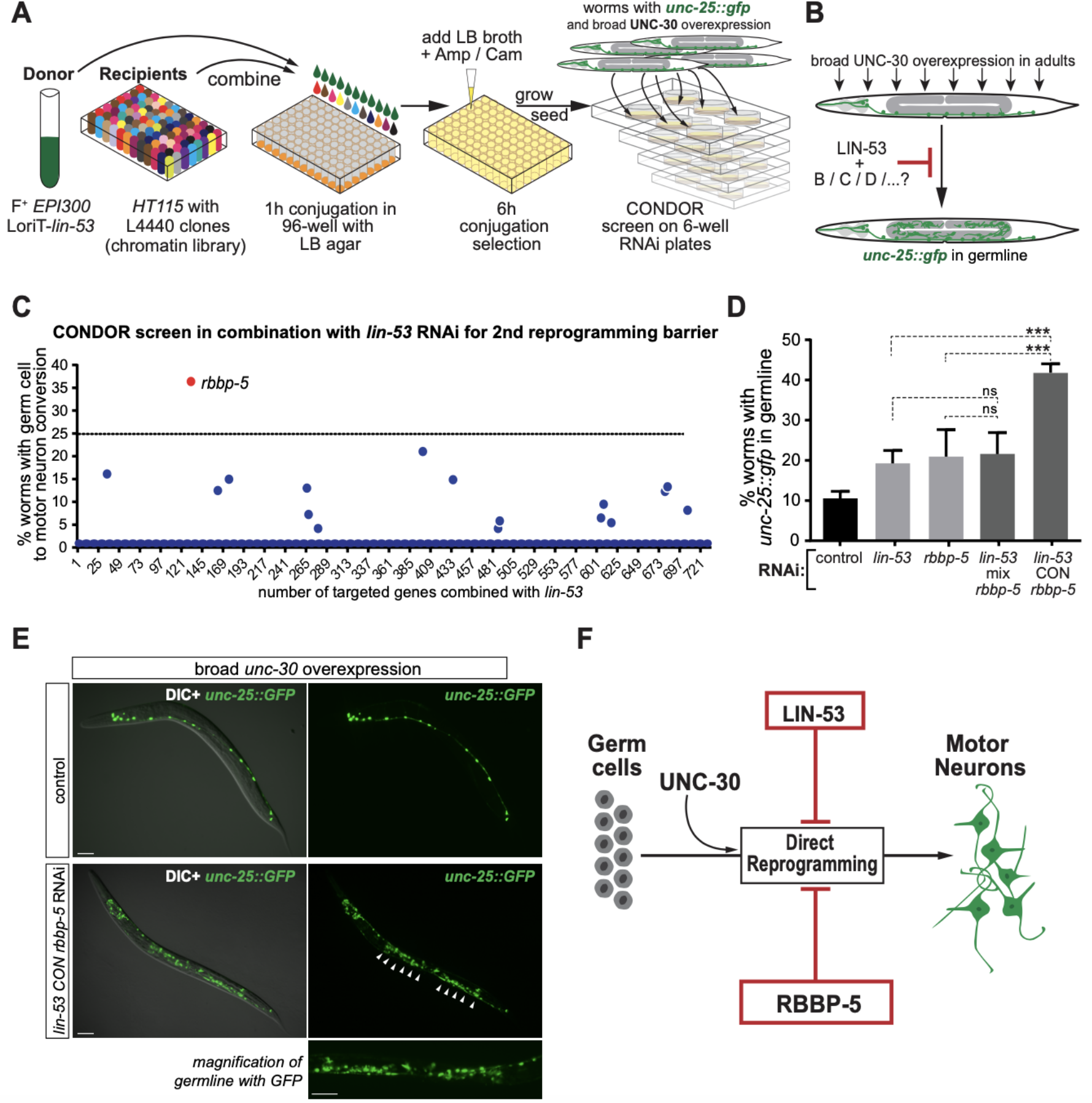
CONDOR identifies RBBP-5 as a novel barrier of germ cell to neuron reprogramming. (A) Illustration of CONDOR to generate double RNAi bacteria targeting *lin-53* together with around 700 other chromatin regulators. F^+^ *EPI300* bacteria containing *LoriT-lin-53* conjugation with *HTT115* RNAi bacteria clones from the previously published chromatin RNAi library (Hajduskova et al., 2019). (B) Transgenic animals expressing the GABAergic motor neuron fate marker *unc-25::GFP* and allowing heat-shock-inducible broad UNC-30 overexpression were fed with conjugated bacteria to assess germ cell to neuron conversion. (C) CONDOR screening for animals with GABAergic fate marker *unc-25::GFP* ectopic expression in the germline. Feeding of *LoriT-lin-53* conjugated with *rbbp-5* RNAi bacteria resulted in a marked increase of *unc-25::GFP* expression induction in the germline. The stippled line indicates the chosen cut-off for enhancement. (D) Direct comparison of *lin-53_CON_rbbp-5* to single RNAi against *rbbp-5, lin-53*, and *L4440-lin-53* mixed with *L4440-rbbp-5 HT115* RNAi bacteria. CONDOR is allowing GABAergic fate reporter expression in the germline more efficiently than mixing RNAi bacteria. Statistics: unpaired t-test; ***p < 0,0001; ns = not significant; Error bars represent SEM. (E) Representative DIC / GFP signal pictures of animals with germ cells expressing the GABAergic neuron fate reporter *unc-25::GFP* upon simultaneous depletion of LIN-53 and RBBP-5 by CONDOR and broad overexpression of UNC-30. White arrow heads indicate germline with reprogrammed cells (this area is shown in magnification below). (F) Model of preventing germ cell conversion to GABAergic motor neurons by RBBP-5 and LIN-53.

Worms were subjected to CONDOR in an F1 RNAi screen and heat-shocked as young adults to induce broad expression of the GABA motor neuron fate-inducing TF UNC-30 (Fig. 6A-B). One day later we scored for animals showing ectopic *unc-25::GFP* expression in their germline (Fig. 6C). Among the more than 700 tested *lin-53* co-depletions, we found that worms fed with bacteria derived from conjugating *LoriT-lin-53* with *L4440-rbbp-5 (lin-53_CON_rbbp-5*) showed a marked increase of the *unc-25::GFP* induction rate in the germline (Fig. 6C). The *rbbp-5* gene encodes for the Set1/MLL methyltransferase complex member RBBP-5 and is the ortholog of human RBBP5 (RB binding protein 5) (Beurton et al., 2019). Next, we compared *lin-53_CON_rbbp-5* to single RNAi against *rbbp-5, lin-53 or L4440-lin-53 mixed with L4440-rbbp-5* RNAi bacteria (Fig. 6D). We noticed that knockdown of *rbbp-5* alone provides a similar number of animals with germlines positive for *unc-25::GFP* as RNAi against *lin-53* (Fig. 6D). However, CONDOR-mediated knockdown of *lin-53* and *rbbp-5* simultaneously (*lin-53_CON_rbbp-5*) almost doubled the efficiencies of single RNAi knockdowns (Fig. 6D). Notably, double RNAi by mixing bacteria yields a similar number of animals with germlines positive for *unc-25::GFP* as single RNAi against *lin-53* or *rbbp-5* (Fig. 6D). A lack of additive or synergistic enhancement could be interpreted as a cooperative function of LIN-53 and RBBP-5 in the same pathway or complex to counteract germ cell conversion. However, CONDOR against *lin-53* and *rbbp-5* indicates an additive effect compared to single RNAi against *lin-53* and *rbbp-5*, rather suggesting functions in separate processes. Together with previous comparisons of double RNAi by mixing versus CONDOR our results provide evidence that CONDOR is suitable for double RNAi to assess genetic interactions, which may be more accurate compared to mixing RNAi bacteria clones.

Overall, we show that our newly developed CONDOR technique is highly efficient to conduct high-throughput double RNAi screens. By using CONDOR we identified the Set1/MLL methyltransferase complex member RBBP-5 as a previously undiscovered germ cell reprogramming barrier demonstrating versatility and robustness of this novel double RNAi technique.

## DISCUSSION

Depletion of gene activities by RNAi-mediated knockdown is essential for investigating gene functions. In particular, genes that cause embryonic lethality or developmental arrest when fully eliminated by knockout approaches (e.g. by mutagenesis or CRISPR/Cas9-mediated excision) can often not be interrogated for their implication in post-developmental processes. Yet, RNAi-mediated co-depletion of such essential genes to investigate complex biological processes in *C. elegans* are challenging. Previous approaches, such as feeding mixed RNAi bacteria, suffer from robustness, as we demonstrate also in this study, or are not practicable for high-throughput screens. For instance, ‘stitching’ together two target genes by cloning into the *L4440* plasmid to produce dsRNA against both targets in the same bacterial cell has been shown to mediate robust double RNAi (Gouda et al., 2010). While this approach is certainly a reliable method for double RNAi, generating, e.g., 700 new plasmids each containing two target genes with subsequent bacterial transformations limits practicability and flexibility for high-throughput double RNAi screens.

We developed a new method of conducting combinatorial RNAi in *C. elegans* based on bacterial conjugation, which we termed CONDOR. By creating a two-component system consisting of conjugation-competent F^+^ *E. coli* (based on *EPI300*), which contain the F-plasmid *pRK24-KanR*, and a modified RNAi donor plasmid *LoriT*, we are able to co-deplete genes simultaneously in a robust manner. We demonstrate CONDOR for model targets (*GFP* and *RFP* in muscle nuclei) as well for several endogenous genes. Sterility due to defective oocyte maturation is caused when both *oma-1* and *oma-2* genes are co-depleted (Detwiler et al., 2001). This ‘*oma*’ phenotype is caused with significantly higher efficiency by CONDOR when compared to mixing the two *oma-1* and *oma-2 HT115* RNAi bacteria. Moreover, feeding conjugated bacteria versus mixed bacteria consistently induced double RNAi with higher efficiency against all tested gene combinations such as *mex-3* and *gld-1* (germline teratomas) as well as *rpn-10* and *rpn-12* (lethality).

Based on the robustness of CONDOR we were able to identify the Set1/MLL methyltransferase complex member RBBP-5 (Beurton et al., 2019) as a novel germ cell reprogramming barrier. The rational for using CONDOR to screen for chromatin factors that counteract germ cell reprogramming was based on our observation that overexpression of UNC-30 in animals with RNAi against the previously identified germ cell reprogramming barrier LIN-53 (Patel et al., 2012; Seelk et al., 2016; Tursun et al., 2011) yielded only a limited number of animals with GABAergic motor neuron fate in germ cells. Feeding worms with conjugated bacteria containing *lin-53* RNAi (*LoriT-lin-53*) and *rbbp-5* RNAi (*L4440-rbbp-5*) led to increased ectopic induction of the GABAergic neuron fate in germ cells.

Notably, comparison of *lin-53_CON_rbbp-5* to single or mixed RNAi against *rbbp-5, lin-53 or L4440-lin-53* mixed with *L4440-rbbp-5* RNAi bacteria revealed almost a doubling of the number of animals with germlines positive for the GABA neuron fate reporter *unc-25::GFP*. In contrast, mixed double RNAi against *lin-53* and *rbbp-5* showed similar numbers as single RNAi against *lin-53* or *rbbp-5*. Such lack of enhancement during genetic interaction testing is usually being interpreted as LIN-53 and RBBP-5 functioning in the same pathway or complex. Generally, it should be mentioned here that RNAi is not ideal to examine genetic interactions. To assess synergistic, synthetic, or additive effects upon loss of two genes, principally the use of null-mutants allow more consistent conclusions. Yet, knockdown by CONDOR of *lin-53* and *rbbp-5* revealed an additive effect compared to mixed double RNAi. This result suggests functions of the chromatin-regulating factors LIN-53 and RBBP-5 in separate regulatory pathways. This notion is supported by previous studies showing that LIN-53 cooperates with the PRC2 chromatin silencer to safeguard the germ cell fate and counteract conversion to neurons (Patel et al., 2012; Seelk et al., 2016). Depletion of LIN-53 or PRC2 subunits resulted in a global loss of chromatin silencing in the germline as revealed by abolished H3K27 methylation (Patel et al., 2012). In contrast, it was demonstrated that RNAi against *rbbp-5*, which is part of the chromatin-regulating complex SET1/MLL/COMPASS (Beurton et al., 2019; Li and Kelly, 2011), reduces H3K4 methylation in the germline. Thus, the enhancement observed upon simultaneous RNAi knockdown of *lin-53* together with *rbbp-5* is likely due to distinct effects on germline chromatin. The finding that LIN-53 and RBBP-5 may act in parallel pathways due to the observed additive effect suggests that CONDOR provides a reliable technique for double RNAi to assess genetic interactions. Mutants of these essential genes can otherwise not be tested in the context of germline safeguarding in adult animals due to lethality or early developmental arrest (Li and Kelly, 2011; Lu and Horvitz, 1998). Generally, double RNAi by CONDOR may provide more accurate genetic interaction testing of essential genes as compared to mixing RNAi bacteria clones.

Yet, the exact mechanism of how RBBP-5 safeguards the germline to prevent conversion to GABAergic motor neuron-like cells remains to be determined and will be subject of future research efforts. Here, we used the identification of RBBP-5 to highlight the power and versatility of CONDOR and provide evidence for its efficiency. As most biological processes are co-regulated by the orchestrated activity of several genes, CONDOR opens new perspectives for all research fields that make use of the genetic model *C. elegans* to address open question *in vivo*. Moreover, robust triple RNAi could be performed by ‘stitching’ two target genes into *LoriT* with subsequent conjugation to other RNAi clones thereby further increasing the multiplexing of knockdowns. Overall, further developments such as CONDOR are likely to increase the complexity of RNAi screens for investigating biological processes in an unprecedented manner.

## MATERIAL AND METHODS

### Nematode cultures

Animals were maintained according to standard procedures (Stiernagle, 2006). Heat-shock sensitive strains were kept at 15°C.

### Caenorhabditis elegans (C. elegans) worm strains

N2: wild isolate, Bristol variant.
BAT28: *otIs305[hsp-16.2p::che-1::3xHA, rol-6(su1006)] ntIs1[gcy-5p::gfp, lin-15(+)] V*
BAT256: *otIs355 [rab-3::NLS::TagRFP] IV*
BAT684: *juIs8 [unc-25::GFP]; barEx147 [hsp-16.2/4::unc-30]*
BAT1616: *ccls4251 [myo-3p::NLS::gfp] I; barIs112 [myo-3p::NLS::tagRFP, HygR] X;*

### Synchronized worm population

Synchronized worms were obtained by bleaching hermaphrodites with eggs or by L1 arrest. Gravid hermaphrodites were treated with household bleach (5% sodium hypochlorite) mixed with 1M NaOH and water (3:2:5) Following worm lysis, eggs were washed three times with M9 buffer. For harvesting L1 worms, plates with freshly hatched L1 larvae were collected by washing off with M9 buffer + gelatin. Arrested L1 larvae and bleached eggs were either applied directly onto RNAi or regular NGM plates.

### Escherichia coli (E. coli) bacterial strains

*OP50*: uracil auxotroph
*HT115*: F-, mcrA, mcrB, IN(rrnD-rrnE)1, rnc14::Tn10(DE3 lysogen: lacUV5 promoter-T7 polymerase) (IPTG-inducible T7 polymerase) (RNAse III minus).
*EPI300*: F-mcrA Δ(mrr-hsdRMS-mcrBC) Φ80dlacZΔM15 ΔlacX74 recA1 endA1 araD139 *SW105: SW103* ΔgalK

### Generation of pRK24-KanR and LoriT

pRK24-KanR (dBT847 Tursun lab name) was constructed by recombineering to replace Ampicillin resistance (AmpR) of *pRK24* (Meyer et al., 1979) with Kanamycin resistance (KanR) because *L4440* used in the standard ‘Ahringer’ *C. elegans* RNAi library (Kamath et al., 2003) already carries AmpR. Recombineering was performed as previously described (Tursun et al., 2009). Primer to PRC amplify the KanR cassette for recombineering were:

FWD: GAA GTT TTA AAT CAA TCT AAA GTA TAT ATG AGT AA ACT TGG TCT GAC AGt tat tag aaa aat tca tcc agc aga cg;
REV: TGT ATT TAG AAA AAT AAA CAA ATA GG GGT TCC GCG CAC ATT TCC CCGAAA AGc gcg gaa ccc cta ttt gt tta ttt ttc.

Generating the ‘donor’ RNAi plasmid *LoriT*, required addition of the *oriT* which allows transfer by conjugation. AmpR of L4440 was replaced with Chloramphenicol (CamR) resistance to allow selection for presence of *LoriT* together with the resident AmpR-containing *L4440* RNAi plasmid after conjugation. Primers used to PCR amplify *oriT* and *CamR* for GIBSON cloning were:

oritFWD: cca ccg gtt cca tgg GGC GCT CGG TCT TGC CTT;
oritREV: cca cgc gtc acg tgg AGC GCT TTT CCG CTG CAT AAC.

Further information can be found in Supplemental Figure 2 B – C and in Suppl. Table 2. *prK24-KanR* and *LoriT* will be made available through Addgene upon publication of this manuscript.

### Generation of donor bacteria: F+ EPI300 with LoriT

The recombination deficient *E*. coli strain *EPI300* is transformed with pRK24-KanR to generate a stable F^+^ strain, which can conjugate with other bacteria. F^+^ *EPI300* (containing *pRK24-KanR*) was made electrocompetent for transformation and aliquots were frozen as previously described (Tursun et al., 2009). Alternatively, F^+^ *EPI300* can be kept as a standard glycerol stocks and made electrocompetent when needed (see below). Gene sequence of the target gene is inserted into the multiple cloning site of the plasmid *LoriT (L4440* plasmid derivative containing *oriT* and *CamR* instead of AmpR). The RNAi plasmid *LoriT-target-gene* is electroporated into F^+^ *E. coli EPI300* to generate the donor RNAi bacteria. In brief, bacteria are grown until reaching an OD600 between 0.4 and 0.8, put on ice for 15 minutes. The cells are pelleted at 4°C for 15 minutes and washed with ice-cold ddH_2_O. The cells are pelleted once more at 4°C and the supernatant is removed except ~0.5 mL. Into an aliquot of 100 μL of F^+^ *EPI300*, 50 ng of donor RNAi plasmid (*LoriT-target-gene*) DNA is being added, transferred to electroporation cuvette (0.2 cm electrode gap) and incubated for 2 minutes on ice. The sample is then electroporated by pulsing with2.5 kV using a standard electroporator. After electroporation 900 μL of LB medium are added and the bacteria are incubated at 37°C for 1 hour under shaking. The cells are plated on selective LB-KanR/CamR plates and incubated overnight at 37°C. Single colonies are picked and grown to prepare glycerol stocks of F^+^ *EPI300* bacteria that are competent for conjugation and can transfer *LoriT-target-gene*.

### Generation of double RNAi bacteria clones by conjugation

On day one 96-well plates filled with 100 μL LB agar are prepared and dried overnight. Donor F^+^ *EPI300* (with e.g. *LoriT-lin-53* as used in this study) and recipient F^-^ *HT115* bacteria of the Ahringer RNAi library clones were grown overnight at 37°C to saturation. Depending on the number of conjugations, the F^-^ *HT115* bacteria with Ahringer RNAi clones should be grown in a 96-well format (deep-well). Donor F^+^ bacteria are pelleted at 4°C and 80% of supernatant is removed resulting in an approximately 5x up-concentration. 5 μL of donor and recipient bacteria are pipetted in equal amounts of on LB agar containing 96-well plates, covered with a lid or aluminum foil and incubated for 1 h at 37°C. After incubation, 100 μL of LB-Amp/Tet/Cam medium is added and the plates are incubated for another 6 hours under mild shaking. Afterwards, about 5 μL from each well are transferred into a new 96 deep-well plate containing LB-Amp/Cam. The second selection step does not contain tetracycline (Tet) as this affects the worms negatively. The 96-deep-well plates are incubated overnight at 37°C under shaking and the conjugated RNAi bacteria are ready to be seeded onto NGM RNAi plates. For further information of RNAi clones used in the study see Suppl. Table 1.

### Determining and confirming successful conjugation (for proof of concept in this study only)

Conjugated RNAi plasmid-containing bacteria clones were confirmed by colony PCR upon growth on Amp/Cam. Donor RNAi plasmid containing bacteria (e.g. *LoriT-lin-53* as used in this study) in the F-plasmid *pRK24-KanR*-containing *EPI300* bacteria were grown overnight to saturation, mixed at a 1:5 ratio on LB agar plates, incubated for 1 hour at 37°C, washed off and plated on LB-Kan/Amp/Cam plates to select for conjugated bacteria clones containing the donor RNAi plasmid and the F-plasmid *pRK24-KanR*.

### RNA interference experiments

The reprogramming experiments were carried out by feeding the animals with bacteria containing conjugated or standard RNAi clones. Generally, we performed F1 RNAi by exposing L4 stage larval animals to RNAi plates. For germ cell reprogramming experiments, plates were kept at 15°C and the following F1 generation was heat shocked at L3/L4 stage for 30 minutes at 37°C. Afterwards animals were kept at 25°C overnight and scored 24 hours post heat-shock for ectopic induction of *gcy-5::GFP* or *unc-25::GFP*. For double RNAi by mixing, bacteria were grown as saturated cultures. The OD600 was measured to ensure that bacteria were mixed at an equal ratio and seeded on standard RNAi plates (Hajduskova et al., 2019). For further information on media recipes used in the study see Suppl. Table 3.

### Microscopy

For imaging, worms were mounted on freshly made 2% agarose pads using 10 mM tetramizole hydrochloride in M9 buffer to anesthetize animals. Microscopy analyses were performed using the ZEISS Axio Imager.M2 (Zeiss) equipped with the Sensicam CCD camero by PCO Imaging. For image acquisition MicroManager plugin in ImageJ was used (Edelstein et al., 2010, 2014). Acquired picture were processed using ImageJ.

## ACKNOWLEDGMENTS

We thank Sergej Herzog for technical assistance. We also thank the CGC, supported by the NIH, for providing strains. This work was partly sponsored by the ERC-StG-2014-637530 and the Max Delbrueck Center for Molecular Medicine in the Helmholtz Association. All procedures conducted in this study were approved by the Berlin State Department for Health and Social (LaGeSo). The authors declare that they have no competing interests.

## AVAILABILITY OF DATA AND MATERIALS

*C. elegans* strains generated in this study will be made available through CGC https://cgc.umn.edu). The plasmids *pRK24-KanR* and *LoriT* will be available via Addgene (www.addgene.org).

## AUTHORS’ CONTRIBUTIONS

MK, CFD, AO and BT conducted experiments, analyzed the data, and helped with the experimental design. MK and BT conceptualized and designed the project, and BT finalized the manuscript. All authors read and approved the final manuscript.

## SUPPLEMENTAL MATERIAL

### SUPPLEMENTAL FIGURES

**Supplemental Figure 1:**
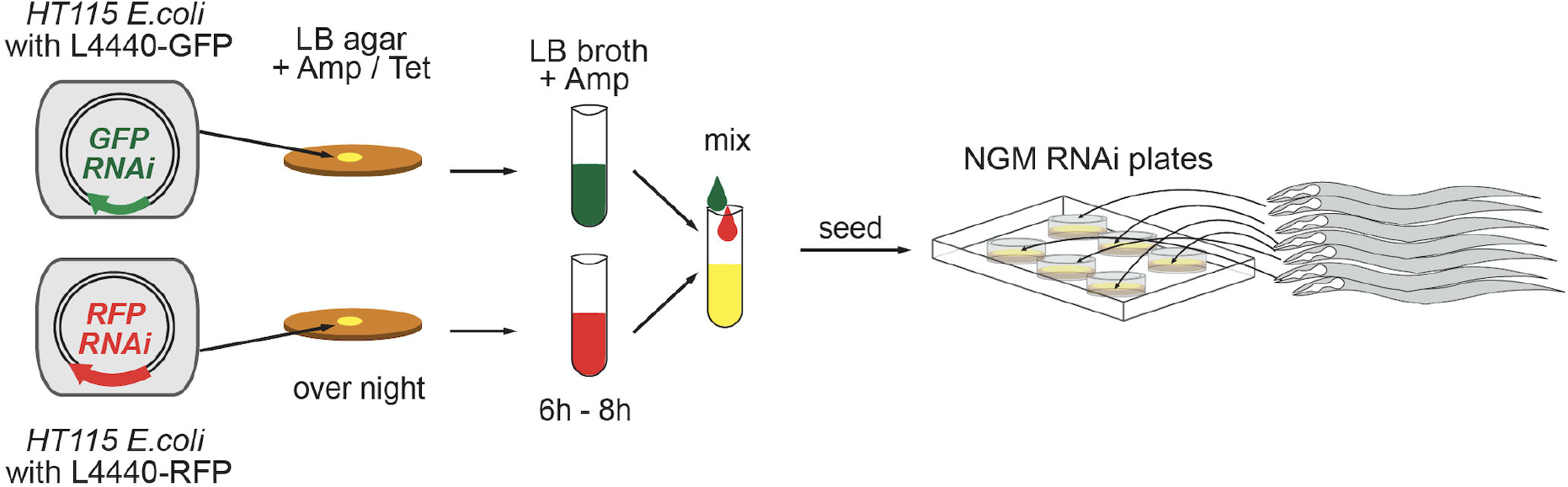
Double RNAi in *C. elegans* by mixing bacteria. RNAi in *C. elegans* is straightforward and can be applied by feeding worms with bacteria that produce dsRNA against the target gene. The standard procedure to perform simultaneous knockdown of two genes is to mix two bacterial strains each producing specific dsRNAs. The illustration shows mixing of bacteria that produce dsRNA against *GFP* or *RFP*. The dsRNA is produced by *HT115 E.coli* bacteria that contain the RNAi plasmid L4440 plasmid. The gene of interest is cloned into L4440, which allows IPTG-induced dsRNA production.

**Supplemental Figure 2:**
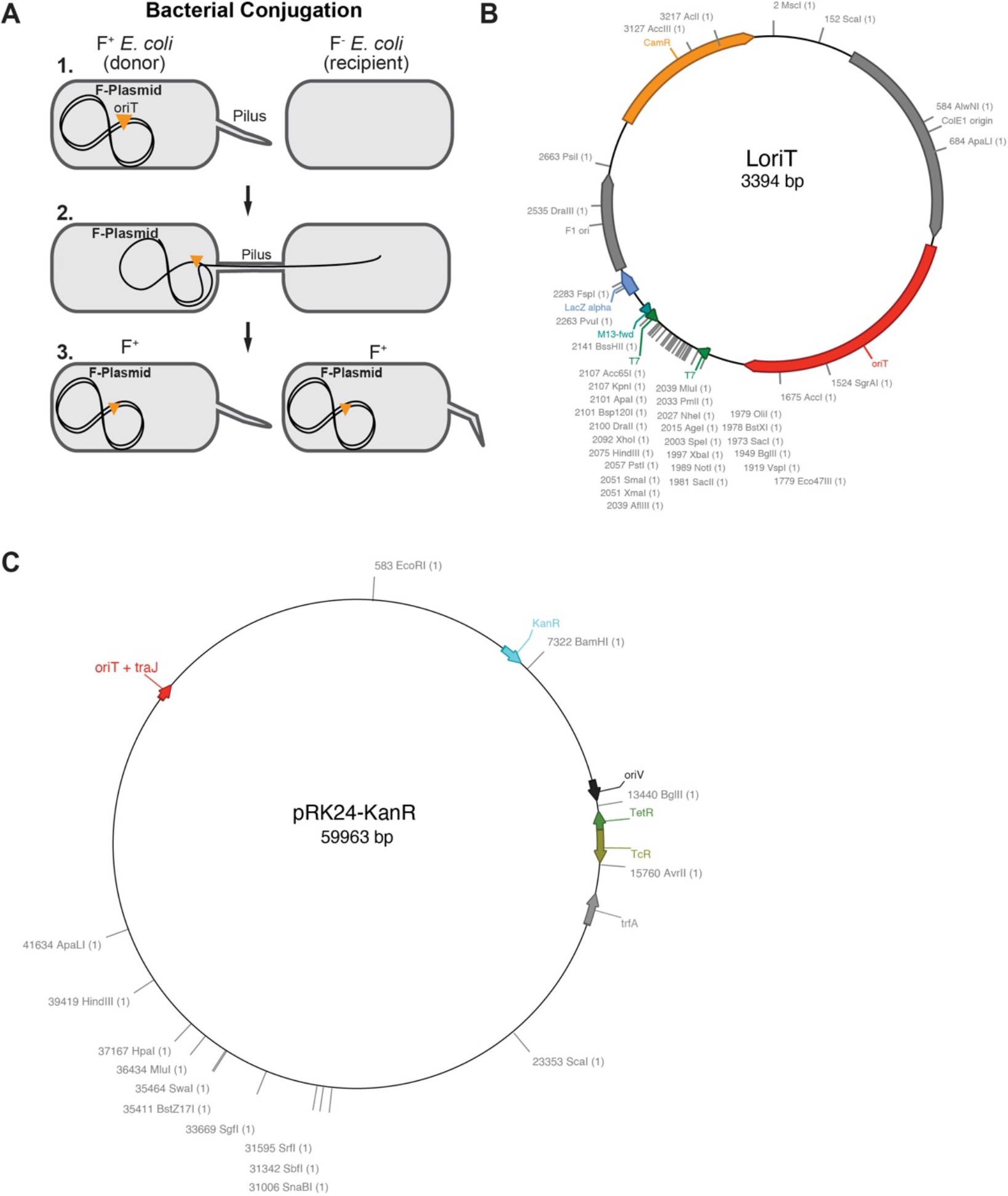
Generating a bacterial conjugation system to combine RNAi plasmids.Generating a bacterial conjugation system to combine RNAi plasmids. (A) Competence for bacterial conjugation requires presence of the fertility factor, also termed F-plasmid, which contains several genes of the *tra* locus for the formation of a pilus appendage. Bacteria with the F-plasmid are denoted as F^+^ (donor) and connect via the pilus to F^-^ bacteria (recipient) to transfer plasmids or other genetic material containing an *oriT* to the recipient. (B) Generating the selectable ‘donor’ RNAi plasmid based on *L4440*, which can be transferred by conjugation, needed the addition of the *oriT* and replacement of AmpR with Chloramphenicol (CamR) resistance. This allows selection for presence of the transferred RNAi plasmid together with the resident AmpR-containing *L4440* RNAi plasmid after conjugation. We termed the newly generated donor plasmid ‘*LoriT*’, which is basically L4440 carrying *oriT* and CamR instead AmpR. (C) To adopt bacterial conjugation for combining RNAi plasmids, we tooke the F-plasmid *pRK24* (Meyer et al., 1979) and replaced the Ampicillin resistance (AmpR) of *pRK24* with Kanamycin resistance (KanR) since *L4440* used in the standard ‘Ahringer’ *C. elegans* RNAi library (Kamath et al., 2003) already carries AmpR. To exchange AmpR with KanR we used recombineering, as previously described (Tursun et al., 2009) due to the extensive size of *pRK24*.

**Supplemental Figure 3:**
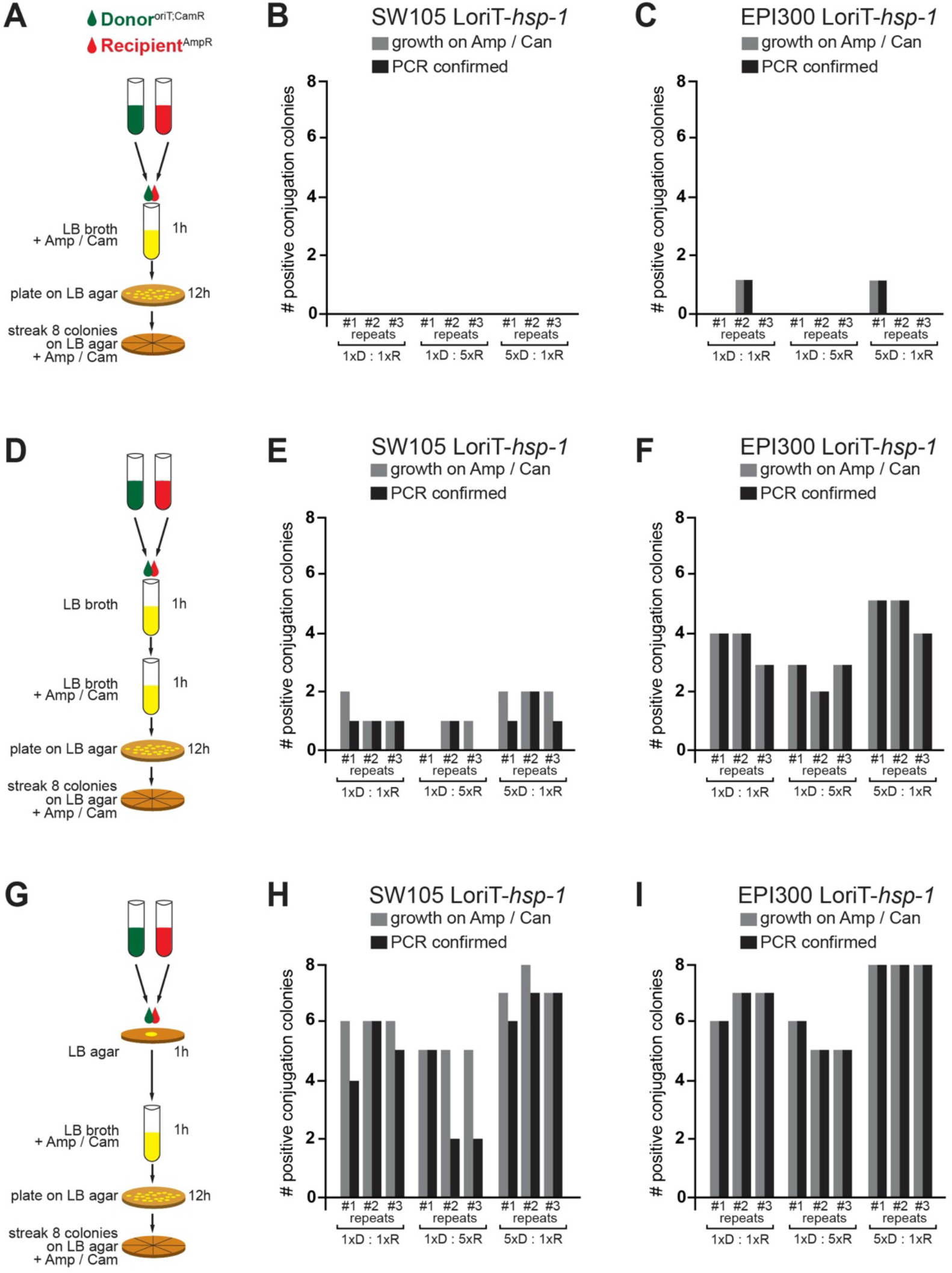
Assessment of conjugation procedures for efficient transfer. (A) Conjugation in liquid culture by combining F^+^ donor bacteria (*SW105* or *EPI300* containing *pRK24-Kan*) and recipient *HT115*. Incubation of donor and recipient bacteria in liquid LB media containing Amp/Cam for 1h and subsequent plating on LB-Agar plates for 12h. The last step of streaking 8 colonies (if any grown) was to test for colony PCR to verify presence of donor (*LoriT-hsp-1*, CamR) and recipient (*L4440-ogt-1*, AmpR). (B and C) The two *E.coli* strains *SW105* or *EPI300* were used previously to handle large DNA constructs such as fosmids (Tursun et al., 2009) and therefore chosen as the host strains for the *pRK24-KanR* episome (F-plasmid for conjugation competence). We aimed for testing 8 colonies from each procedure of conjugation either combing a ratio of 1:1, 1:5, or 5:1 of donor D and recipient R bacteria. In some cases no colonies were obtained. Obtained colonies were tested by PCR to confirm successful conjugation. The procedure as shown in (A) performed overall poorly. (D) Conjugation by combining donor and recipient bacteria in liquid LB media without antibiotics for 1h, and then with Amp/Cam for 1h with subsequent plating on LB-agar plates to select at least 8 colonies (if any grown) for examining by PCR. (E and F) As for (B and C) but with more obtained colonies. Still the yield is low and *SW105* F^+^ donor bacteria appeared to perform very poorly. As before, we could not even obtain 8 colonies for this procedure as shown in (D) to test. (G) Conjugation on solid LB-agar without antibiotics by combining donor and recipient bacteria for 1h. Afterwards, incubation in liquid LB broth with Amp/Cam for 1h (either directly adding liquid LB, if performed in 96-well or transferring colony to culture tube) with subsequent plating on LB-Agar plates (Amp/Cam) to select at least 8 colonies for examining by PCR. (H and I) this procedure yielded the most efficient conjugations, however the use of *SW105*-based donor bacteria showed less robustness. The use of 5:1 (donor D: recipient R) yielded highly efficient conjugation with correct conjugation in all tested cases.

**Supplemental Figure 4:**
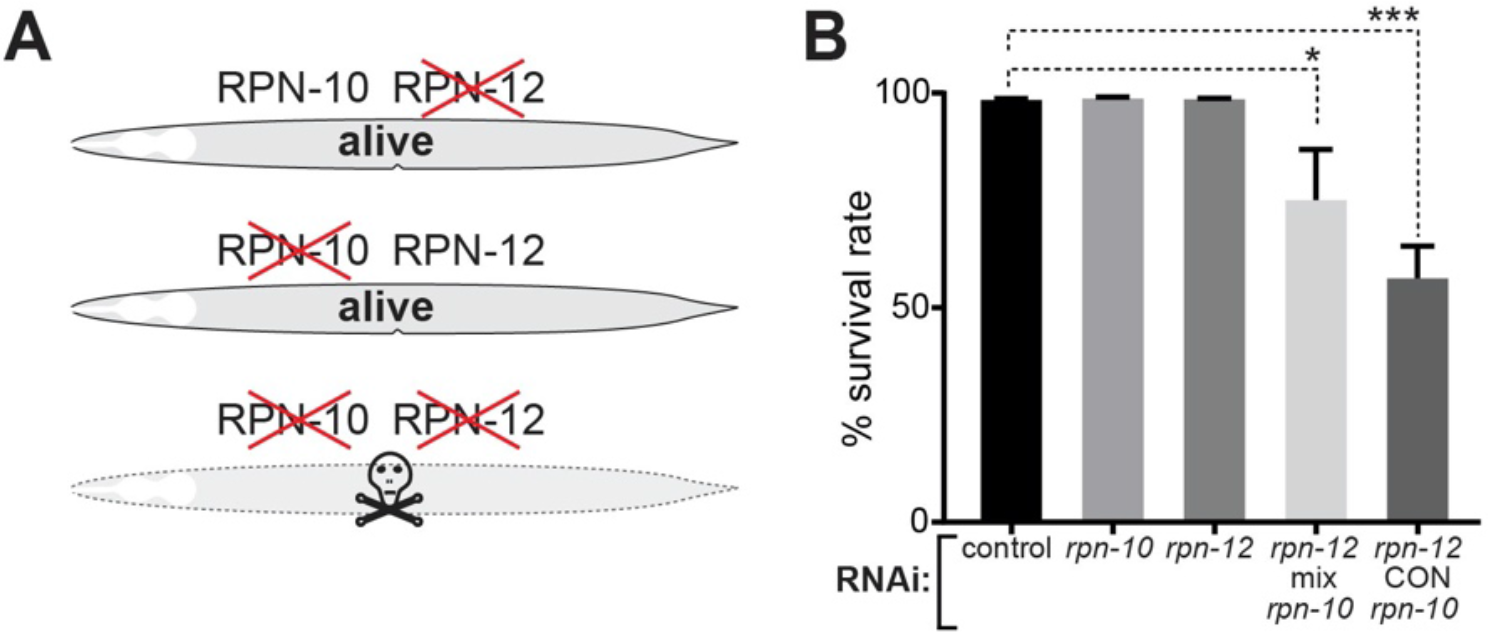
Synthetic lethality induced upon co-depletion of proteasomal subunits. (A) We targeted the 26S-Proteasome subunit genes *rpn-10* and *rpn-12*, which cause synthetic lethality when co-depleted (Takahashi et al., 2002). (B) CONDOR-mediated simultaneous knockdown of *rpn-10* and *rpn-12* reduced survival by around 50%. In contrast, 25% of the animals fed with mixed *rpn-10* and *rpn-12* RNAi bacteria died indicating that CONDOR is more efficiently depleting *rpn-10* and *rpn-12* simultaneously.

### SUPPLEMENTAL TABLES

**Supplemental Table 1:** RNAi clones used in the study

| Target Gene | Function | Source |
| --- | --- | --- |
| *lin-53* | histone-chaperone LIN-53 ortholog of human RBBP4 | Chromatin RNAi library; Hajduskjova et al., 2019 |
| *rpn-10* | proteasome subunit | Chromatin RNAi library; Hajduskjova et al., 2019 |
| *rpn-12* | proteasome subunit | Chromatin RNAi library; Hajduskjova et al., 2019 |
| *rbbp-5* | Retino blastoma protein binding Protein; Set1/MLL methyltransferase complex member | Chromatin RNAi library; Hajduskjova et al., 2019 |
| *oma-2* | Oocyte Maturation defective; Zn-Finger | Ahringer RNAi library; Kamath et al., 2003 |
| *gld-1* | Translational regular; ortholog of human QKI - KH domain containing RNA binding | Ahringer RNAi library; Kamath et al., 2003 |
| *mex-3* | ortholog of human MEX3A RNA binding family member | Ahringer RNAi library; Kamath et al., 2003 |
| *ogt-1* | ortholog of human OGT (O-linked N-acetylglucosamine (GlcNAc) transferase | Chromatin RNAi library; Hajduskjova et al., 2019 |

**Supplemental Table 2:**
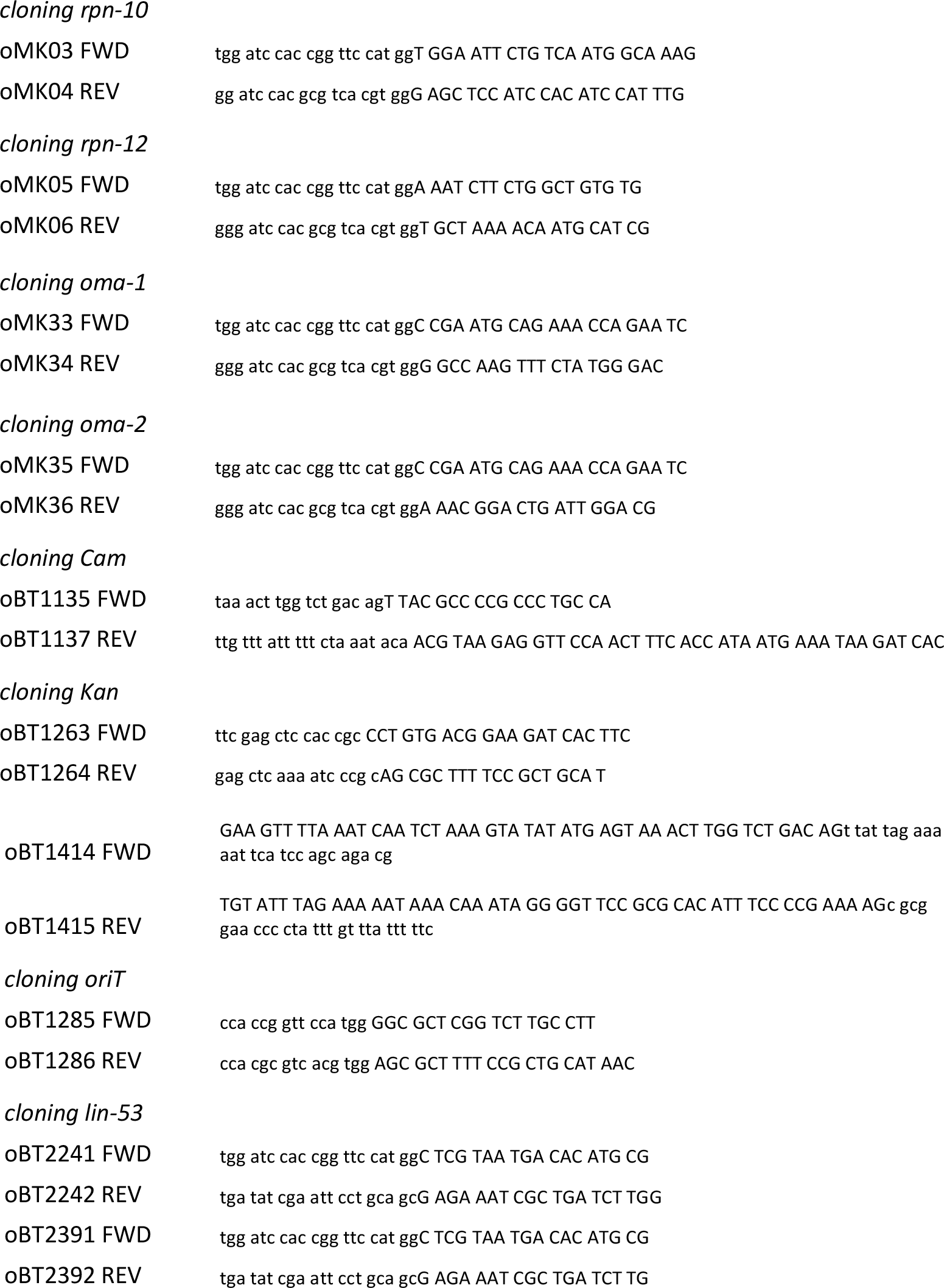
Primers used in the study

**Supplemental Table 3:**
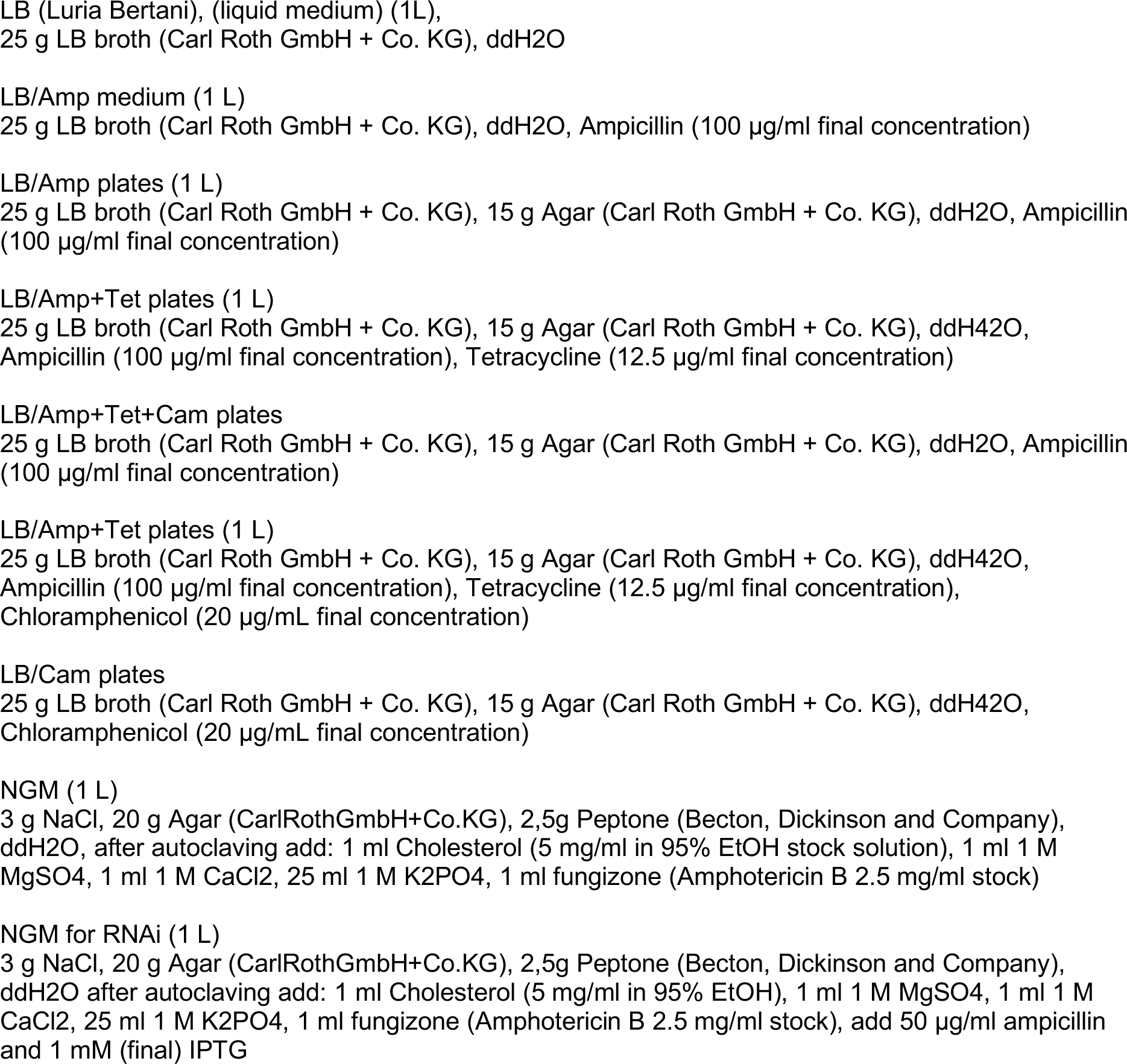
Media recipes used in the study

